# On the evolution of emergence patterns driven by environmental cycles

**DOI:** 10.64898/2026.10.07.757275

**Authors:** Dharanish Rajendra, Jule Neumann, Tobias S. Kaiser, Chaitanya S. Gokhale

**Affiliations:** Chair for Computational and Theoretical Biology, Julius-Maximilians University, Klara-Oppenheimer-Weg 32, Würzburg, 97070, Germany; Max Planck Research Group “Biological Clocks”, Max Planck Institute for Evolutionary Biology, August-Thienemann-Str. 2, Plön, 24306, Germany; Joint Institute for Individualisation in a Changing Environment (JICE), University of Münster and Bielefeld University, Germany

**Keywords:** lunar rhythms, phenotypic variability, replicator-mutator equation

## Abstract

Regular environmental cycles such as day and night or the lunar cycle are ubiquitous and drive many biological processes. Many organisms have evolved endogenous time-keeping mechanisms to anticipate these cycles to accurately time their behavior or synchronize physiological processes. The evolution of such synchronization to environmental cycles and their variability is not very well understood. As an example, we use the marine insect *Clunio* (Diptera, Chironomidae), which synchronize adult emergence and reproduction with the lunar cycle, and show different emergence patterns across populations. We use a model constructed with replicator-mutator dynamics to provide a novel outlook at the evolution of these emergence patterns. In particular, we investigate the effect of two traits of individual phenotypic variability of emergence, the spread and shape, on population-wide emergence patterns. We find that the semilunar and lunar emergence patterns are alternatively stable and occur when the phenotypic distribution has a narrow and is roughly flat. Arrhythmic emergence occurs when the phenotypic distribution is wide and nearly flat. With our model, we introduce the concepts of frequency-dependent selection and individual phenotypic variability into the theory of chronobiology, which gives us insights into the evolution of rhythmicity in these organisms.

## 1. Introduction

Environmental cycles such as the seasons and the day-night cycle influence various biological rhythms. Plants and trees shed their leaves during autumn and bloom during spring. Mammals, e.g. bears and rodents, can enter a dormant state of hibernation during winters. Many species of birds migrate towards the equator before winter and return to their breeding grounds before the summer. While some of these phenomena are direct responses to changing external stimuli, many rhythms are controlled by an internal biological clock, which anticipates the changes in the environment and allows the organism to prepare in advance. Circadian clocks are well studied Patke, Young and Axelrod (2020) (Nobel Prize for Physiology or Medicine 2017) and a paradigmatic example of a circadian rhythm is the sleep-wake cycle. In contrast, lunar and semilunar rhythms are understudied, even though they are found across the animal and plant kingdoms and are important for reproduction (Kaiser and Neumann, 2021; Numata and Helm, 2014; Ritter and Tessmar-Raible, 2024; Ben-Attia, Reinberg, Smolensky, Gadacha, Khedaier, Sani, Touitou and Boughamni, 2016; Kronfeld-Schor, Dominoni, de la Iglesia, Levy, Herzog, Dayan and Helfrich-Forster, 2013). One of the most famous examples of a reproduction event synchronized to the lunar cycle is the annual spawning of the Palolo worm (*Eunice viridis*) in the Pacific Ocean. More than 70 years of documentation of this event at the Samoan islands emphasized the precision with which those worms time their reproduction to the third lunar quarter (Naylor, 2010). However, the existence of a truly endogenous circalunar clock has only been experimentally verified in a few species (Kaiser and Neumann, 2021). We choose one of them, the marine insect *Clunio*, as the subject for our study. *Clunio* is a marine genus of non-biting midges of the family Chironomidae (Diptera). They inhabit the intertidal zone of coasts across the globe (Neumann, 1976). The intertidal zone is that region of the coast which experiences cycles of submergence and exposure due to the tides with varying water levels over the lunar cycle. *Clunio*’s reproduction is tightly synchronized with the lunar cycle (Fig. 1A).

**Figure 1:**
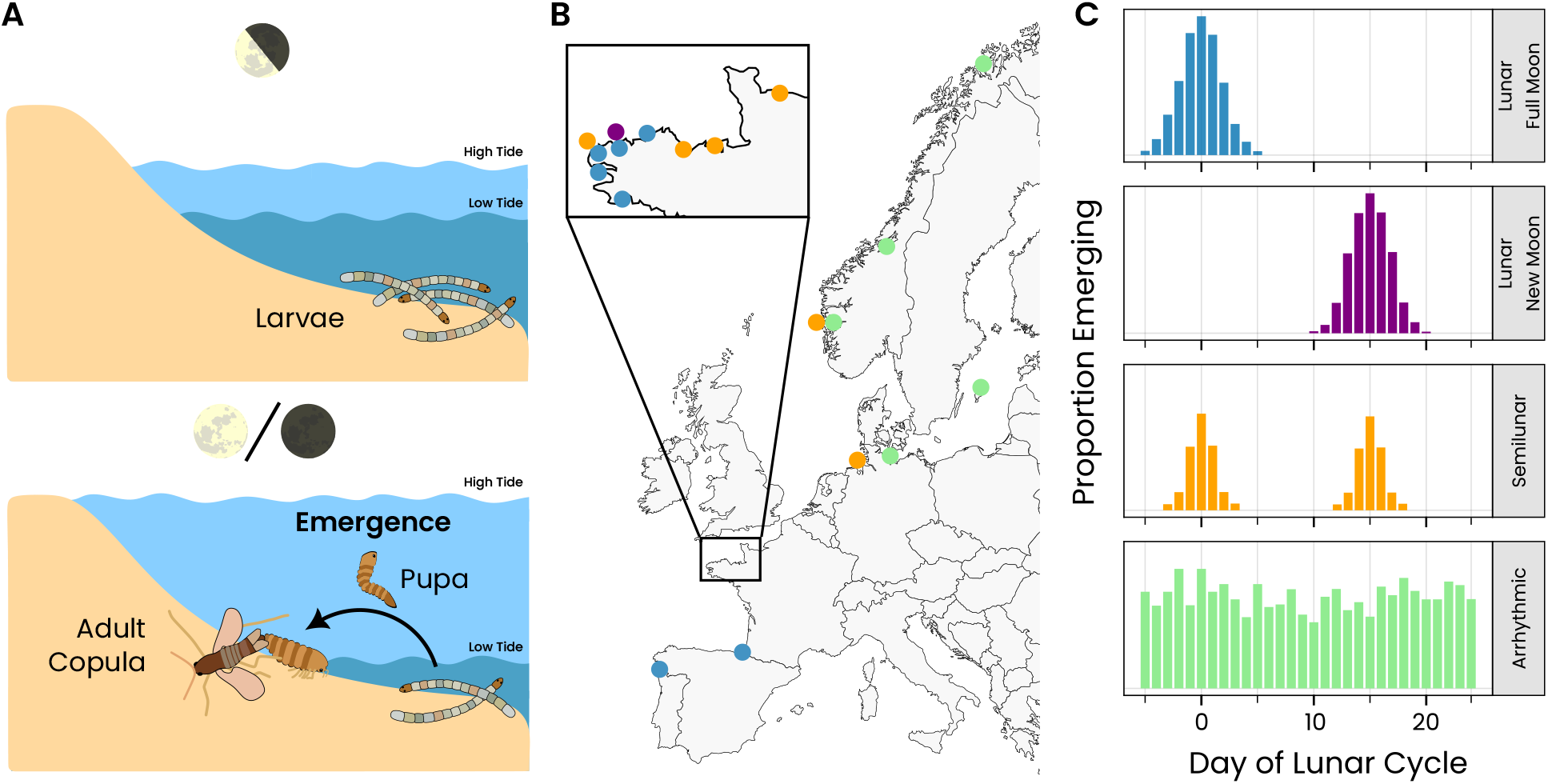
**A**. The life cycle of the marine midge *Clunio* is synchronized with the lunar cycle. Adults emerge only during the spring tides at full and new moon when most of the intertidal zone is exposed, and mate and lay eggs on the exposed surface. **B**. *Clunio marinus* populations across the coasts of Europe display distinct emergence patterns. **C**. These emergence patterns include lunar (emerging at full or new moon), semilunar (emerging around full and new moon) and arrhythmic. Each emergence pattern is given a distinct and consistent color across B & C and also in the subsequent figures. These plots are an idealized interpretation of the experimental data reviewed in Neumann and Kaiser (2023)

**Figure 2:**
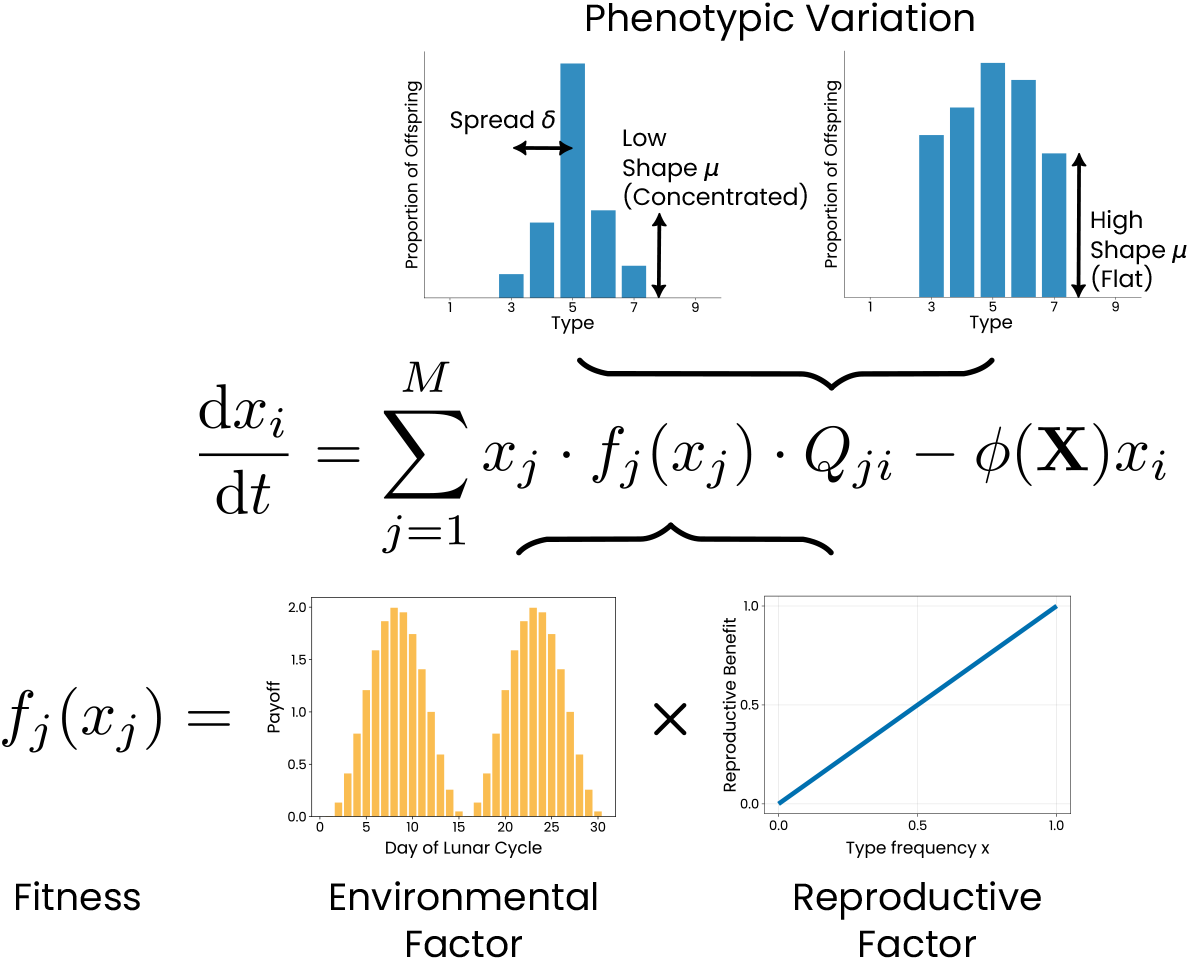
Overview of the components of the model. The central equation of our model is the replicator-mutator equation. There are *M* of these equations for *i* = 1, …, *M*, corresponding to each type of individual (this is the day of emergence over the lunar cycle), where the dynamical quantities are the frequencies of the different types of individuals *x*_*i*_. The left-hand side is the instantaneous rate of change of the frequency of type *i* in the population. The first term on the right is the effective fitness of type *i*, which is calculated by averaging the fitness of neighboring types, taking into account the phenotypic variability. The fitness of an individual is the product of the environmental factor, which takes into account the tides, and the reproductive factor, that takes into account the frequency of the focal type. The phenotypic variability goes into the *Q*-matrix, whose rows represent the distribution of offspring of a single parent across different days, and is controlled by the parameters of spread *δ* and shape *µ*. The population average fitness is denoted by *ϕ*.

After hatching from an egg, the midges spent most of their lives (a few months) underwater in the larval stage (Neumann, 1976). After a brief pupal stage, they eclose and emerge as adults. The adult stage lasts only a few hours during which they mate and lay eggs onto the exposed substrate. Adult emergence and, thus, reproduction is highly synchronized with the lunar cycle. At spring tides, which occur during full and new moon, the amplitude between the low and high tide is the maximum. Reproduction around the spring low tides ensures that oviposition occurs in the lower intertidal zone, which prevents larvae that hatch from these eggs from drying out.

Populations of *C. marinus* across different geographic locations display a diversity of emergence patterns (Fig 1 B,C). On the coast of southern Europe, populations exhibit a lunar emergence pattern. That is, they have a single emergence peak per lunar cycle, either around the full moon or around the new moon. Towards the north, in addition to these, one can find semilunar strains which display two emergence peaks per lunar cycle, around both full and new moon. Further to the north, are arrhythmic strains which, unlike the other two, do not have any lunar periodicity and display a uniform emergence pattern throughout the lunar cycle (Neumann and Kaiser, 2023). The populations along the coast of Norway emerge daily synchronized just to the tidal cycle as daylight is not a reliable cue in the arctic and the populations at the Baltic Sea emerge with a circadian rhythm due to weak tides and deposit their eggs into the water (Honegger and Neumann, 1969). Note that, the lifespan of an individual midge is, in general, longer than a single lunar cycle. To explain the above-mentioned population level differences in emergence patterns, we investigate the impact of individual variability. By individual variability, we mean the variation of the offspring emergence time. This can arise from phenotypic plasticity as well as genetic variability. The distribution of individuals with different phenotypes then results in a population level phenotype.

Most theoretical work in chronobiology so far has focused on modeling the endogenous oscillator itself (Gonze, 2011; Forger, 2017; Roenneberg, Chua, Bernardo and Mendoza, 2008; Tyson, Albert, Goldbeter, Ruoff and Sible, 2008; Abraham, Granada, Westermark, Heine, Kramer and Herzel, 2010; del Olmo, Legewie, Brunner, Höfer, Kramer, Blüthgen and Herzel, 2024; Becker-Weimann, Wolf, Herzel and Kramer, 2004). These works study the molecular and cellular mechanisms of the endogenous oscillator and how it reacts to external stimuli including modeling the gene-protein interaction networks, cell-cycle progression and signal processing. This work derives a lot from the theory of (coupled) oscillators, which itself has been studied extensively (Chandra and Scott, 1983). Daido (2002) and Rand, Shulgin, Salazar and Millar (2006) do consider evolution, but it is in the context of circadian rhythms. A circalunar clock, on the other hand, can be based on a completely different mechanism and functional principle and can also depend on the functioning of the endogenous circadian clock (Neumann, Rajendra and Kaiser, 2024). Thus, theoretical studies on circalunar rhythms are lacking, especially within the context of evolution and phenotypic variability.

To fill this gap, we use a model based on replicator-mutator dynamics (Nowak, 2006), that is agnostic to the mechanism of the internal clock. The “replicator” part captures the process of selection and controls the fractions of different variants in the population based on their fitness, which can be frequency-dependent. The “mutator” part captures the generation of variation, genetic and non-genetic. The overall equations then balances selection with variation. It can tell us the trajectory of a population given some initial conditions or the end-point of dynamics after a sufficient amount of time. In the following section Model & Results, we develop the model part by part and also describe the results in tandem. We start off with a description of the general model before tailoring the parameters to the lunar strains of *Clunio*. Following which, we relax an assumption about frequency dependent mating to bring the model closer to reality. Next, we make an inclusion of new types of individuals with different phenotypic variability into the model to capture the semilunar strains as well. This model shows that all four emergence patterns of *Clunio* occur when the distribution of phenotypic variation is flat. However, the arrhythmic strain occurs only when the spread of the distribution is high, while lunar and semilunar strains occur when it is low. The semilunar and two lunar strains are alternatively stable, meaning that they can exist in the same environmental conditions and phenotypic variability, but not coexist. Finally, we discuss the implications of our results, how they can be generalized to other species and also comment on why studying the evolution of these processes is powerful.

## 2. Model & Results

We consider an infinite population of individuals which exist in an environment with cycling conditions of a period of *M* days. Correspondingly, there are *M* types of individuals in the population which emerge on each of the *M* days of the environmental cycle. We do not assume that the lifecycle of the individuals has the same duration as the environment cycle. The individuals undergo a developmental period and when they are sufficiently developed, emerge on the day corresponding to their type in the upcoming cycle. We assume that all the individuals have the same developmental period but are only separated in phase (day of emergence). The proportions of the different types of individuals are *x*_*i*_, *i* = 1, …, *M*. The fitness of each type depends, in general, on the whole composition of the population.

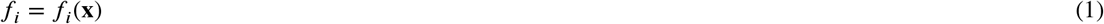

Since the adult lifespan of *Clunio* is just a few hours, individuals can only mate with another individual of the same type. That is, they need to emerge at the same phase *M*. We assume that physically searching for a mate is the rate limiting step in reproduction. Thus, we say that the fitness of a type is proportional to just its own frequency in the population. That is, the more there is of a type, the easier it is for them to find a mate.

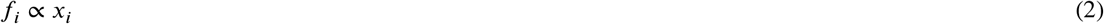

In addition to this, the lunar day on which a parent emerges and produces offspring affects the survival of the larvae and hence the fitness of the type. This comes from the understanding that larvae need to be submerged, and their survival rate is related to the proportion of time spent underwater. Larvae that are not constantly submerged are likely to dry out and die. The proportion of time being submerged depends on the depth of the larvae in the intertidal zone. After adult emergence and mating, eggs are laid close to the water level. As a simplification, we assume the depth of the eggs, and hence the larvae, is the water level at the time of emergence. This proportion of time submerged can be approximated by the absolute cosine function

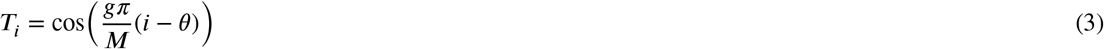

Where, *g* is a constant to make sure the tides have the correct frequency over the *M* days, and *θ* is the phase of the tides. Since the strength of tides varies across locations, a parameter was introduced to capture it. Thus, the environmental factor in the fitness is defined as follows

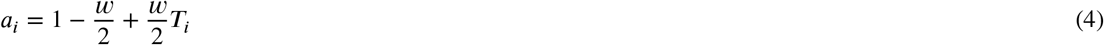

where, *w* is the strength of selection, capturing the strength of the tides. Putting these two factors together, the fitness of type *i* is:

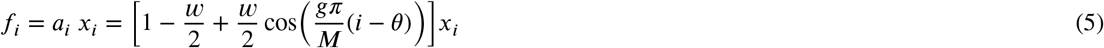

Further, individuals display variability, i.e. offsprings do not necessarily emerge on the same day as their parents. Mathematically, this is captured by a *M* × *M* sized matrix *Q* that determines the offspring phenotype.. The element *Q*_*ij*_ of the matrix denotes the proportion of offspring of parent type *i* that will emerge on the *j*th day. We consider two traits/parameters that control the phenotypic variability in the *Q*-matrix: the phenotypic variation spread *δ* and the shape *µ. δ* controls the width of the distribution of phenotypic variability i.e. the maximum distance (in time) from the parent that the offspring can emerge. A larger *δ* means that offspring have a possibility to emerge farther in time from the day of emergence of the parent. *µ* controls the shape of this distribution. Low *µ* means that the distribution is concentrated at the center while high *µ* means the distribution is more spread out or flat. In other words, a higher *µ* implies a higher probability of an offspring to emerge away in time from the parent, within the allowed spread.

The specific form of *Q*-matrix that we use is banded and treats each parent equally. It has the form:

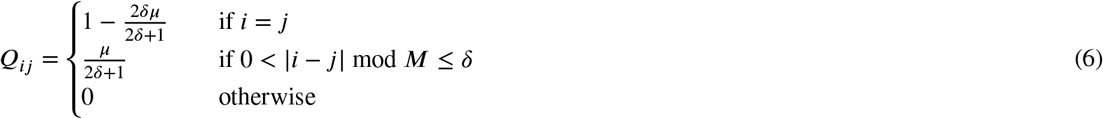

If the offspring emerges on the same day as the parent, then *i* = *j* and the probability of that happening is 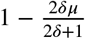. Offspring can emerge on either of the *δ* days before or after this day. The probability of the offspring emerging on of these days is identical and is 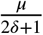. When *µ* = 0, there is no phenotypic variability and all offspring emerge on the same day as the parent. When *µ* = 1, all 2*δ* + 1 days have equal probability for emergence.

With these fitnesses and *Q*-matrix, we can write down the replicator-mutator equation that governs its dynamics (Nowak, 2006):

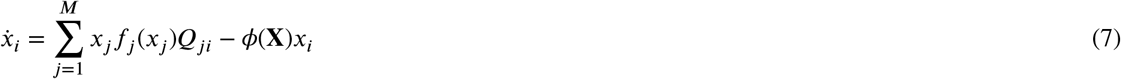

Where, *f*_*i*_(*x*_*i*_) is the fitness of type *i* as in Eq. 5 and 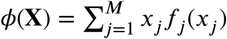 is the average population fitness.

We are interested in the equilibrium or steady-state solutio^*j*=^n^1^s to this set of equations. We obtain these solutions by setting all derivatives on the left-hand side of these equations to zero 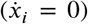 and solving the remaining set of non-linear equations. That is, we find sets of *x*_*i*_s that solve the equations. A solution to this set of equations represents a whole population. Multiple solutions i.e. populations cannot coexist. However, a single solution can contain one or more coexisting timing strains.

We make a note that phenotypic variability and the *Q*-matrix are essential to the model. Without it, we only get solutions where all individuals emerge on the same day of the lunar cycle. Analytical calculations supporting this are shown in SI section S1: Without variability.

### Lunar and Arrhythmic Strains

In the case of the lunar cycle, we consider the number of days in the cycle to be 30 (approximated from the actual 29.53 days of a lunar cycle), and set *M* = 30. We numerically solve the above replicator-mutator equation in the equilibrium case for a fixed value of selection strength *w* and a wide range of values of phenotypic shape parameter *µ* and phenotypic spread *δ*. The effect of changing *w* is presented in SI section S3: Effect of selection strength. The number of distinct stable equilibria for each parameter combination are shown in Figure 3A. When the shape parameter is low, there are 30 unique, stable emergence patterns or strains, corresponding to emergence on each of the 30 days of the cycle. These strains do not coexist but rather exist exclusively. Due to the low phenotypic shape parameter, the probability of offspring emerging on very different days from the parent is very low. Here, frequency dependence in selection plays a much stronger role than the environmental fitness effect. Due to this, populations starting off emerging on non-optimal days (according to the environmental fitness benefit), are stuck there and are not able to evolve to emergence on a more optimal day. This is why there are 30 stable solutions for all possible days of the cycle. As the shape *µ* is increased and one moves towards the right on this plot, the number of stable solutions decreases gradually. This occurs because the solutions with emergence on the days with the lowest environmental payoff factor become unstable at *µ*. The reason for instability is that the environmental fitness effect gradually rises compared to frequency dependence of reproduction. As populations which are emerging on non-optimal days are not completely stuck and can now evolve to emerging on a more optimal day. Thus, as the *µ* increases, solutions with low environmental fitness start disappearing from the set of possibilities.

**Figure 3:**
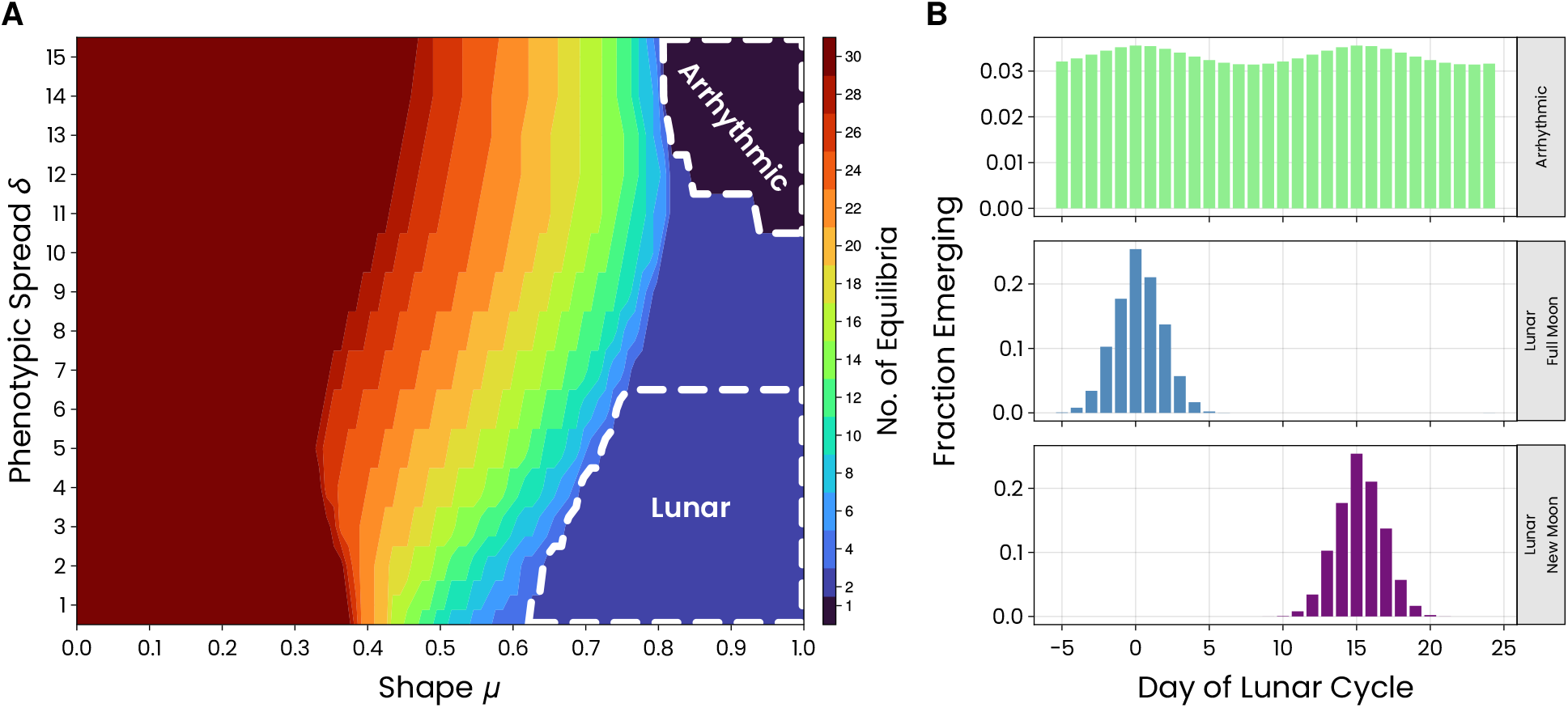
**A**. Number of stable equilibrium solutions of the equations with *M* = 30. The regions of the *µ*–*δ* parameter space having a different number of solutions are indicated using colors. The dark-brown region on the left contains 30 distinct solutions or strains, one for each day of the cycle. In this region, shape parameter is very low, meaning that large changes from parent to offspring are not possible. This results in the effect of frequency dependence of reproduction being dominant. This prevents populations starting off on a non-optimal day from evolving towards emerging on the optimal day. Moving from the left to the right, the number of stable solutions decreases. The blue region on the bottom-right has only 2 distinct solutions, and the dark blue region on the top-right has only one. The dashed lines mark the regions with the named strains. The region of the arrhythmic strains has only one viable solution and has roughly equal emergence on all days of the cycle including the highest tides. The region of the lunar strains has a single peak per lunar cycle, concentrated around full moon or new moon. In between these two regions, we see intermediate solutions which have non-zero but unequal emergence on both tidal peaks. **B**. Example stable equilibrium solutions of the equations from the highlighted regions in the parameter space in A for the arrhythmic strain and the two lunar strains. The colors for the emergence patterns depict the different strains as shown in Fig. 1. Other solutions are depicted in Fig. S2.

A majority of this parameter space does not correspond closely to any observed population or strain in nature (see S2), with the exception of the equilibrium solutions with a high shape parameter (Fig. 3B). For a high shape parameter, the number of stable solutions depends on the phenotypic spread as well. When both shape and spread are high, the arrhythmic population is the only stable solution. But when spread is low instead, there are two stable solutions. These are the lunar full moon and new moon strains. Both are alternatively stable, meaning that they can exist under the same conditions. However, they cannot coexist. All three solutions are shown in Figure 3B. In the wild, however, the full moon and new moon strain can coexist. This implies the existence of additional mechanisms not considered yet or the violation of assumptions of our model.

### Non-linearity of Frequency-Dependent Mating

We previously assumed that the act of finding a mate is one of the biggest factors affecting fitness. That is, we assumed strong frequency dependence in mating with a proportional relationship between reproductive payoff of an individual and the frequency of its type in the population. However, this relationship does not need to be linear. It could be rapidly increasing for low *x* and the increase can slow down for higher *x*, or the other way around. Thus, to relax this assumption of linearity and to take into account other relevant possibilities as well, we modify the fitness function and introduce a new parameter *h*:

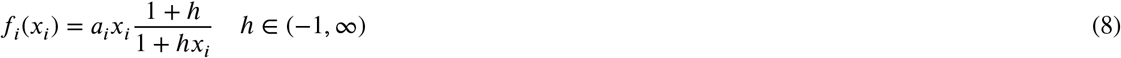

The parameter *h* controls the curvature (concavity or convexity) and non-linearity of the fitness function. For *h* > 0, it is concave and has diminishing marginal benefits. For *h* < 0, the function is convex and is accelerating or has increasing marginal benefits. This kind of non-linearity in frequency-dependence of mating can arise due to factors such as environmental conditions and the way in which *Clunio* physically searches for mates. This curve for different values of *h* is depicted in Figure 4A.

**Figure 4:**
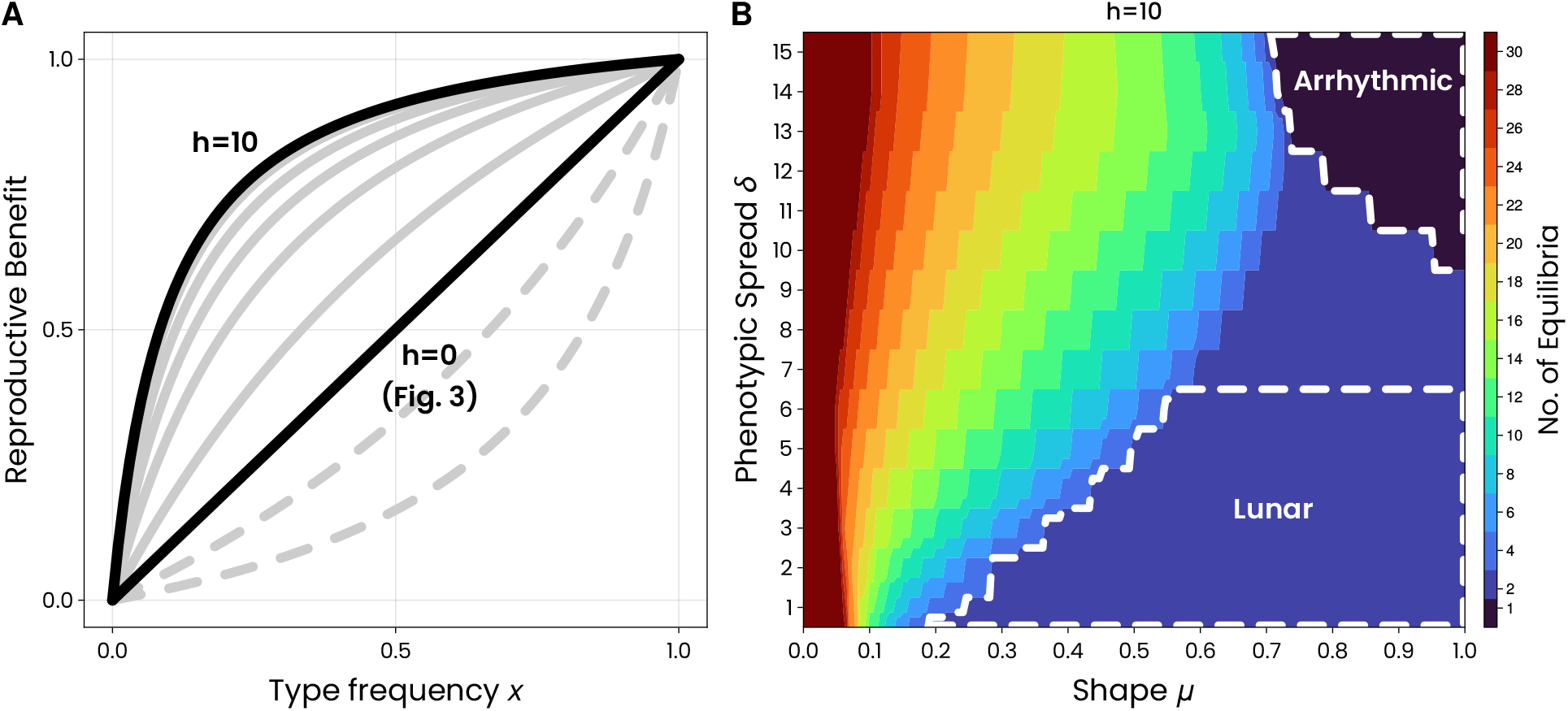
**A**. Reproductive fitness of an individual as a function of the frequency of its own type in the population for different values of *h*, the curvature of frequency dependence. As *h* increases, the curves become more concave and there is a decreasing marginal benefit. The gray solid lines show the curves of reproductive benefit for *h* = 1, 3, 5, 7, 9, with the gray dashes lines corresponding to *h* = −0.5, −0.8. **B**. Number of equilibria of the replicator-mutator equation with *M* = 30 and for high concavity *h* = 10. The regions corresponding to the strains observed in nature are also indicated by the white dashed line. These regions are now larger with higher *h* (compare with Figure 3A).

The effect of high *h* on the equilibria in the whole parameter space is shown in Figure 4B. The main effect of high concavity on reproduction is that the regions of the realistic equilibria (lunar and arrhythmic) increase in size. This means that at a higher population density, lunar and arrhythmic strains are now possible for a wider range of shape parameter. The reverse holds true for when *h* is below zero and the regions with realistic equilibria (lunar and arrhythmic strains) decrease in size. As the concavity increases (*h* > 0), the environmental benefit imposed by the tides now plays a larger role. Thus, equilibria with emergence on days with low environmental benefit become unstable at a lower shape value *µ*. This results in the boundaries separating regions holding different number of equilibria shifting towards lower *µ*, as the curvature *h* is increased. This results in the widening of the region holding the lunar and arrythmic strains.

### Inclusion of Multiple Periodicities

So far, in the model, all types (*i* = 1, …, *M*) have had the same periodicity. That is, they all emerge once every *M* days and *M* is the same for all types. However, in nature, we have different strains with different periodicities. Semilunar strains emerge every 15 days, and lunar strains every 30 days. In order to account for these different emergence patterns, we combine types having different *M* into the same model. Specifically, we have 30 different types with *M* = 30, each of which emerging on a different day of the 30-day cycle. The proportions of these types are denoted by *x*_*i,l*_, *i* = 1, …, 30. In addition, we include 15 different types with *M* = 15, each of which emerge on a different day of the 15-day cycle but twice in the 30-day cycle. The frequencies of these are denoted *x*_*i,sl*_, *i* = 1, …, 15. Thus, we have 45 different types in the population. We make a simplifying assumption that the semilunar and lunar individuals do not mate with each other. This simplification is realistic because these different types generally emerge at different times of the day in nature and around one of the two daily low tides. Due to their short adult lifespans, they may not meet each other (Kaiser, von Haeseler, Tessmar-Raible and Heckel, 2021).

The fitness of the semilunar types is the same as the fitness of the lunar types emerging on the same day, because fitness is only dependent on the tides which is determined by the day of emergence.

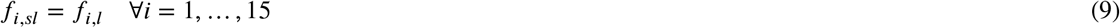

Thus, we have two different *Q*-matrices. *Q*_*l*_, a 30 × 30 matrix for the reproduction of the lunar types and *Q*_*sl*_, a 15 × 15 matrix for the reproduction of the semilunar types. We say that the phenotypic spread of the semilunar types is half that of the lunar types. This is because the maximum possible values themselves are different.

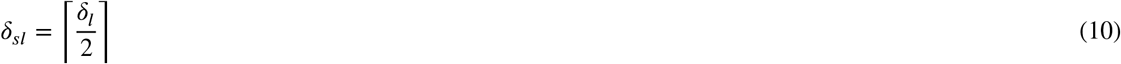

Solving this system of equations including both semilunar and lunar types, and calculating the number of solutions each parameter combination, we get a plot that is qualitatively similar to the one with just lunar types (Fig. 5A, compare with Fig. 3A). We have 45 stable solutions in the left region of the parameter space with low shape *µ*. And as *µ* increases, the number of stable solutions gradually drops off until *µ* reaches a high value. At high *µ*, depending on what value the phenotypic spread takes, the number of solutions can either be 1, 2, 3. The regions with 1 and 2 solutions correspond to the arrhythmic strains as shown in Fig. 5B row 1. The solutions in the arrhythmic region are not completely flat - they have some slight humps and dips. In the region with 2 solutions, the two solutions only differ slightly in the magnitude and phase of these humps, as they arise from either the lunar types or from the semilunar types. However, they can both be still considered as arrhythmic due to their emergence across the lunar cycle. In the region with only 1 strain, the lunar and semilunar types both produce arrhythmic strains. However, they differ less than the tolerance of the numerical solver or 10^−6^. Thus, the two are considered as the same solution. In the lower part of the region with 3 solutions, the solutions correspond to the two lunar strains and the semilunar strains, respectively. The solution of the semilunar strain is distinct from a combination of the lunar full moon and new moon strains together.These 3 solutions are alternatively stable and cannot coexist, similar to the case with only types of periodicity *M* = 30.

**Figure 5:**
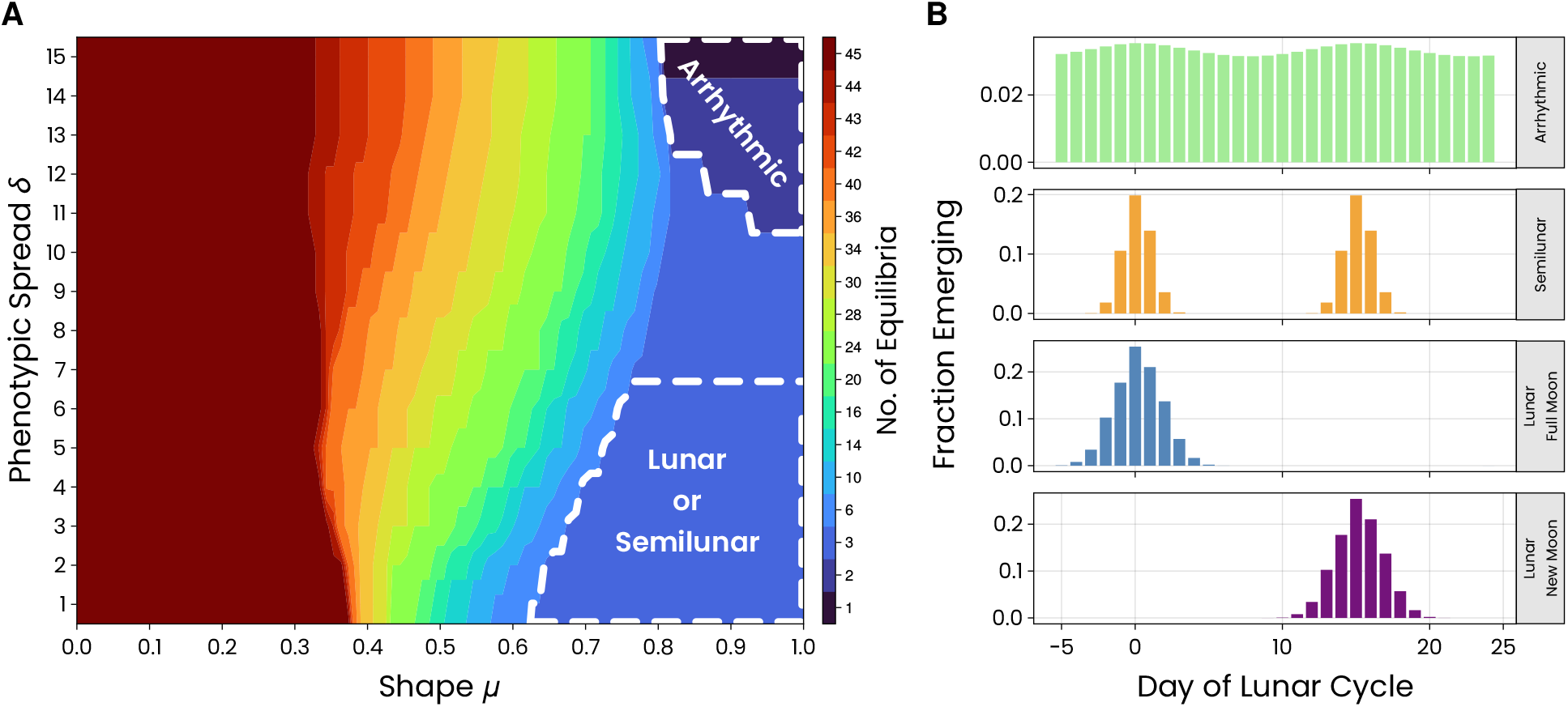
**A**. Number of solutions for the replicator-mutator equation with a combination of types with *M* = 30 and also *M* = 15. The different regions of the parameter space having different solutions are indicated by colors. The two regions showing where arrhythmic strains and semilunar & lunar strains occur are marked by the white dashed line. **B**. Representative solutions from the highlighted regions in the parameter space in A showing the arrhythmic strain, the semilunar strain and the two lunar strains. The colors for the emergence patterns depict the different strains as shown in Fig. 1.

## 3. Discussion & Conclusion

Our theoretical model demonstrates that even under an unchanging fitness landscape, the phenotypic trait variation can determine whether a population is arrhythmic or rhythmic. That is, when the shape parameter is high, increasing spread parameter changes the emergence pattern from lunar or semilunar to arrhythmic. Moreover, all strains observed in nature occur only in a particular region of the parameter space, that is characterized by a high shape parameter or a flat phenotypic variation distribution. Biological or physical constraints may be preventing the shape parameter from being low. The variations in the cues on moonlight and tides, and error in their sensing and in the subsequent synchronization of their biological clock can prevent highly consistent emergence. The solutions with low shape parameter may be of low fitness and hence not observed. Maintaining a low shape parameter may come at a cost, but may not confer the individuals with a sufficient benefit to be maintained. There may also be other unconsidered factors, which confer low fitness or instability to the individuals having a low shape parameter. The three rhythmic strains, full moon, new moon and semilunar, are alternatively stable, meaning that they can occur at the same trait values of phenotypic variation. While the semilunar strain required a new type in the model to be well captured, the full moon and new moon strains can transition from one to another given sufficiently high perturbations or stochasticity. This may suggest that the new moon and full moon lunar strains are evolutionary more closely related to each other than the other strains. However, this could also just be due to the way in which the model is constructed. When increasing the concavity of frequency dependence in reproduction (increasing *h*), the region of the parameter space which shows the lunar, semilunar and arrhythmic strains increases in size. This means that if a population has a high concavity (non-linearity) in frequency dependence, the semilunar, lunar and arrhythmic strains can be captured by a wider range of phenotypic trait parameters. All of this shows that the different strains of *Clunio* can be captured by a single model just by varying a small number of parameters.

Even at the same parameter values of phenotypic variability there could be multiple alternative stable states with different emergence patterns. And the one that is realized at a particular location could depend on the initial conditions and also on stochasticity. The model also predicts that none of these different strains can coexist at the same place and at the same time. However, we do observe coexistence of multiple strains of different period and phase in nature (Kaiser et al., 2021). Therefore, there must be a mechanism in play not considered in the model or some of the assumptions violated. One such mechanism could be the existence and utilization of different niches of some sort. One of the strong assumptions made in the model, for mathematical simplicity and tractability, is that of (infinitely) large population sizes. If a population of the full moon strain and new moon strain do coexist at a point in time, any small difference in population composition will result in a difference of fitness between the two strains because of the frequency dependent fitness. Any small difference will result in the change in population composition in the same direction. Due to the infinite population size, this continues until the strain with higher fitness dominates, and the other goes extinct. A relaxation of this assumption and the stochasticity arising from small and finite population sizes might stabilize the coexistence of multiple strains.

There are a few further simplifications that this model makes. The model assumes that individuals are characterized only by the day of their emergence and a strain by its lunar emergence pattern. However, in nature, an individual is also characterized by the time of emergence during the day. Further, there are more than just one full moon strain or one semilunar strain. In nature, there are multiple strains in different (and even the same) locations showing the same lunar emergence rhythm but differ in other aspects, such as diurnal emergence rhythms, sensitivity towards stimuli or zeitgebers and adaptation towards local moonlight and tidal regimes (Kaiser, 2014). Taking into account these other dimensions of the problem could explain the coexistence. Further investigation into these factors is required to determine the mechanism behind coexistence.

To generalize, the population-level emergence patterns driven by an environmental cycle depend heavily on the individual phenotypic variability. This model and its results are not specific to the system of *Clunio* but could be applied to any organism that shows an emergence or reproduction pattern related to an external environmental or ecological cycle. The main features of *Clunio* that are captured in this model are (a) single emergence and reproduction event per lifetime, (b) a fitness that is dependent on the time of emergence, and (c) a limited window for mating. These features results in a strong selective pressure on the time of emergence, and hence mating. If all of these features are present in another system, then this model can be directly applied to it. Depending on the number of peaks in the payoff landscape, we would get a different number of alternatively stable strains at high shape and low spread parameters. If the adult stage of the organism is longer than the interval between two emergence time-points, then the fitness function *f*_*i*_**(*X*)** should be modified to capture mating between different emergence types. These results show the different solutions and under which trait values they exist and are stable. However, it says nothing about which trait values will be selected for in different environmental conditions. Including an additional evolutionary process that would modify and act on the characteristics of the distribution would be a natural extension to this work, and could provide us insight about possible evolutionary trajectories taken by the strains in nature.

## CRediT authorship contribution statement

**Dharanish Rajendra:** Writing – review & editing, Writing – original draft, Visualization, Methodology, Investigation, Formal analysis, Conceptualization. **Jule Neumann:** Writing – review & editing, Writing – original draft. **Tobias S. Kaiser:** Writing – review & editing. **Chaitanya S. Gokhale:** Writing – review & editing, Writing – original draft, Investigation, Supervision.

## S1. Without Variability

If we ignore any variability in the emergence of an individual, i.e. the offspring emerges on the same day as its parent, we can set the *Q*-matrix to identity *I*. Then, the governing equation reduces to:

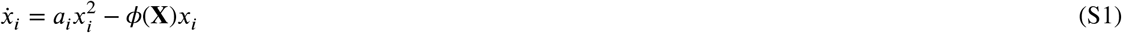

We can solve for the steady state solution by setting 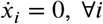. This becomes:

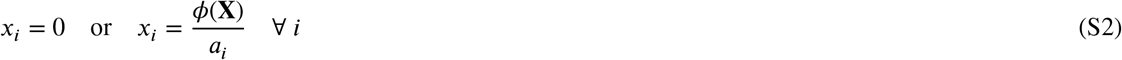

There exists a simple family of localized solutions, *M* in number:

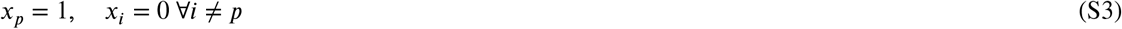

for some *p ∈* 1, …, *M*. In these solutions, all of the individuals emerge on a single day of the cycle. And there are *M* number of solutions corresponding to localized emergence on each day of the cycle.

However, there are several other solutions, with coexistence of any number of types. For coexistence of *K* types of individuals, there are 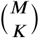 number of solutions. So, the total number of solutions is: 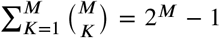. For a value of *K* and set of indices *C* ={*l*_1_, *l*_2_, …, *l*_*K*_} *⊆* {1, 2, …, *M*}, an equilibrium solution is:

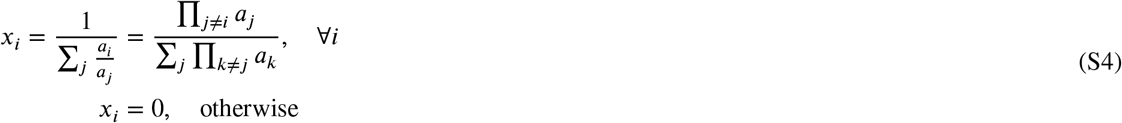

The sums and products are all carried out over the set of indices *C*. For example, a coexistence solution of types *p* and *q* is given by:

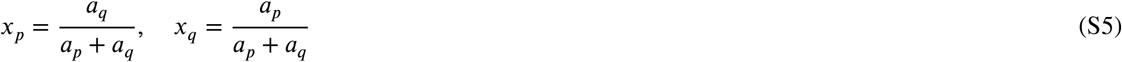

To determine the (linear) stability of these solutions, we have to construct the Jacobian matrix *J* as follows:

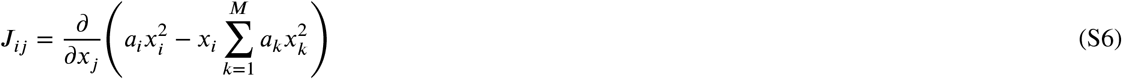

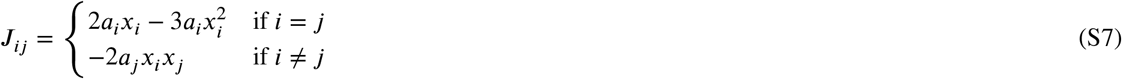

For the family of localized solutions of *x*_*p*_ = 1 for some *p*, the Jacobian is given by:

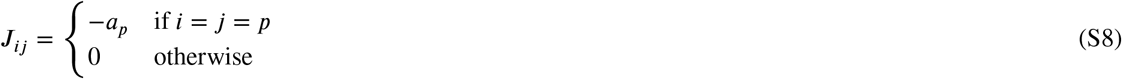

Therefore, these solutions are always locally stable as long as its environmental fitness benefit is positive, *a*_*p*_ > 0.

Let’s now check the stability of the general equilibrium solution 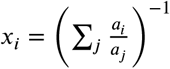. The trace of the Jacobian matrix is equal to the sum of the eigenvalues of the matrix, Tr(*J*) = ∑_*i*_ *λ*_*i*_. The trace is given by:

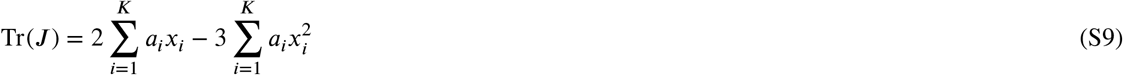

where, *K* is the number of types in the coexistence solution. The type indices are rearranged so that indices in *C* are first followed by the other indices which are not part of the solution. Substituting the general solution in the trace, and simplifying, we get the expression:

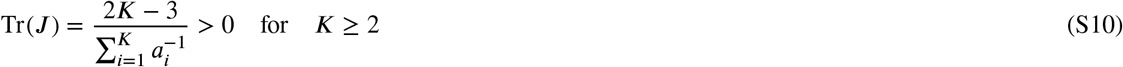

Therefore, solutions involving coexistence of 2 or more types has at least one positive eigenvalue, indicating that they are unstable. Therefore, in this situation, only solutions with localized emergence of all individuals on a single day are possible.

### Basins of Attraction

Each of the localized equilibrium solutions are locally stable. Thus, each of them will have their own basin of attraction. That is, a region in the space of population compositions, and any population starting in this region will end up with that corresponding solution after a long time. The size of the basin of attraction of a solution will tell us how likely that particular solution is likely to be seen if the starting composition is randomly chosen.

Because of the nature of the system, each of the basins of attraction is a convex polyhedron in the M-1 simplex with vertices as the unstable equilibria neighboring the stable equilibrium. Thus, the fraction of volume of the whole state space occupied by the basin of attraction can be computed by computing the volume of these polyhedra. These calculations are done numerically. The total volume of the simplex is 1/(*M* − 1)!. We set the payoff vector to be:

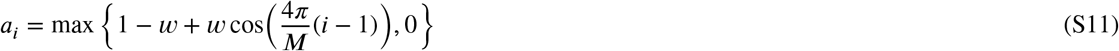

and perform this calculation for a range of values of selection strength *w* for *M* = 7. The calculation of the volume of a convex hull of points is computationally intensive and it becomes infeasible for high dimensional simplexes. Therefore, we perform the calculations for a relatively low value of *M* = 7. We expect the results to be qualitatively similar for higher *M* as well. We plot the relative size of the basin of attraction for each type when *M* = 7 for a range of *w* in Fig. S1. The environmental fitness benefit is just the term cos 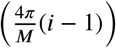 in the payoff vector. We see that when there is no selection, each of the solutions have the same size of basin^*M*^of attraction. That is, each of the solutions are equally likely to occur when initial conditions are randomly chosen. As the strength of selection increases, the types with a higher environmental fitness benefit increase in basin size while the other ones decrease in basin size. Even at high strength of selection, many types having low environmental fitness benefit have a non-negligible basin of attraction. This means that if the initial conditions are chosen at random, there is a significant chance that the system will end up with a population emerging on a sub-optimal day.

### Summary

If there is no individual variation in the emergence times, then the only possible “strains” are with individuals that all emerge on the same day. There are several different such possible strains corresponding to localized emergence on the different days of the lunar cycle. At low selection strength, then each of the stable solutions are equally likely when starting from a random population composition. As the strength of selection increases, the strains consisting of high fitness types gain a relatively larger basin of attraction. As the strength of selection becomes high, some of the strains consisting of low fitness individuals becomes unstable. However, even at high selection strengths, there is more than one possible stable strain.

## S2. Solutions to the equations

In Fig. S2, we show the solutions of the equations for different parameter combinations across the *µ*-*δ* phase space. For a shape of *µ* = 0.3, we are in the region with 30 stable solutions. These solutions have almost the whole population emerging on the same day, with a small proportion emerging on other days. As phenoyptic spread *δ* is increased here, the proportion of individuals emerging on the central day reduces a bit. There are 30 different solutions for this value of the *µ*, all having same distribution, but only differing in shape.

As the shape parameter is increased to *µ* = 0.5, the emergence on the central day decreases further, but it still dominates in the population. For both these values of *µ*, the solutions are qualitatively similar for different phenotypic spread *δ* When the shape parameter is increased past the transitional phase to *µ* = 0.85, the solutions vary highly with *δ*. For low *δ* = 2, we see the lunar strains (full moon or new moon). For high *δ* = 14, we see the arrhythmic strain. As the spread parameter increases slightly from 2 to 4, the solutions still roughly represent the lunar strains, but with emergence on a higher number of days of the lunar cycle. But when *δ* is increased to 9, the solutions no longer represent the lunar strains, but something entirely different.

When the shape parameter is increased further to *µ* = 1, there changes in the solutions are slight. All solutions broaden out a bit. This results in the arrhythmic strain’s emergence pattern becoming more flat, closer to a true arrhythmic strain.

## S3. Effect of Selection Strength

In the main text, we have shown and discussed the phase diagram for a single value of selection strength *w*. The range of possible solutions remains qualitatively the same as *w* changes. However, it has a significant effect on the *µ* − *δ* phase diagram, as seen in Fig. S3. The phase diagram retains the same general characteristics for all values of *w*. That is, towards the left, there is a large region with 30 stable solutions. Each of these solutions correspond to the whole population emerging very tightly around a single day in the lunar cycle, and there are 30 different solutions corresponding to emergence for each day of the cycle. As the shape parameter *µ* is increased beyond a certain value (depending on the spread *δ*), the solution with the lowest fitness (due to low environmental fitness benefit) becomes unstable. As *µ* continues to increase, more and more previously stable solutions become unstable, in order of increasing fitness. At low *δ*, as *µ* is increased beyond an appropriate value, we reach a region with only two stable solution, corresponding to the two lunar strains. And similarly, at high *δ*, as *µ* is increased, we reach a region with only one stable solution, corresponding to an arrhythmic strain.

As the selection strength (*w*) is changed, the sizes and shapes of these different regions in the phase diagram changes. At low *w*, as in Fig. S3A, the region with 30 stable solution is very large. These solutions change continuously as the shape parameter *µ* increases, and the one with lowest fitness becomes unstable at a relatively high value of *µ*. From this point, the number of stable solutions decreases rapidly and reaches 1 or 2 just with a small increase in *µ*. As *w* is increased, as in Fig.S3B,C, the region with 30 stable solutions shrinks. The transitional region where the number of solutions decreases grows in width. That is, the number of solutions changes more gradually with *µ*. However, the region with 1 and 2 solutions (with lunar and arrhythmic solutions) does not change much in size.

As *w* increases, the difference between maximum and minimum environmental benefit increases, and so does the difference between maximum and minimum fitness of solutions. Due to this larger difference in fitness, the lower fitness solutions become unstable at lower values of phenotypic shape parameter. This results in the shrinking of the region with 30 solutions. Because of the increased difference in fitness of the different solutions, the low fitness solutions become unstable at different values of the shape parameter *µ*. This results in the widening of the transitional region. These two factors combine and consequently, the size of the regions with only 1 and 2 solutions remains roughly the same.

**Figure S1:**
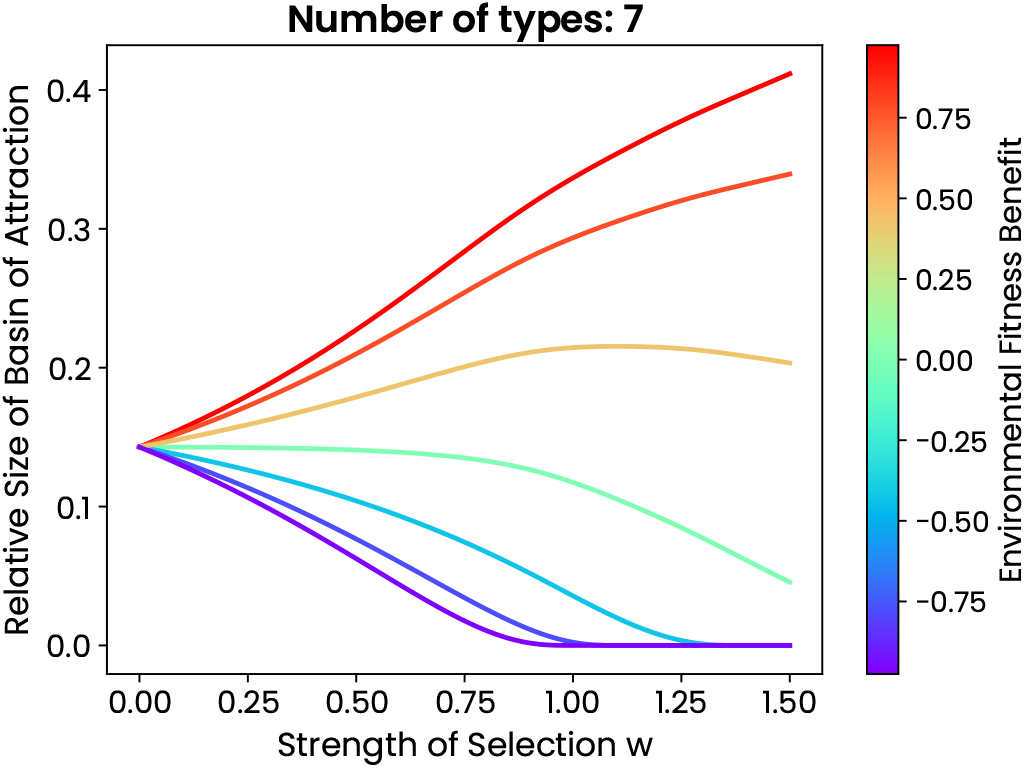
The relative size of the basin of attraction of each type of individual compared to the total size of the state space simplex as a function of the strength of selection *w*. The individual curves are colored by the environmental fitness benefit experienced by them. The size of basins of attraction indicate the following. If one were to randomly assign a initial composition for a population and let it evolve according to these dynamics, what fraction of them will end up conforming to each of the stable solutions. This plot shows that even at high selection strength, there is more than one stable solution with a sizeable basin of attraction.

**Figure S2:**
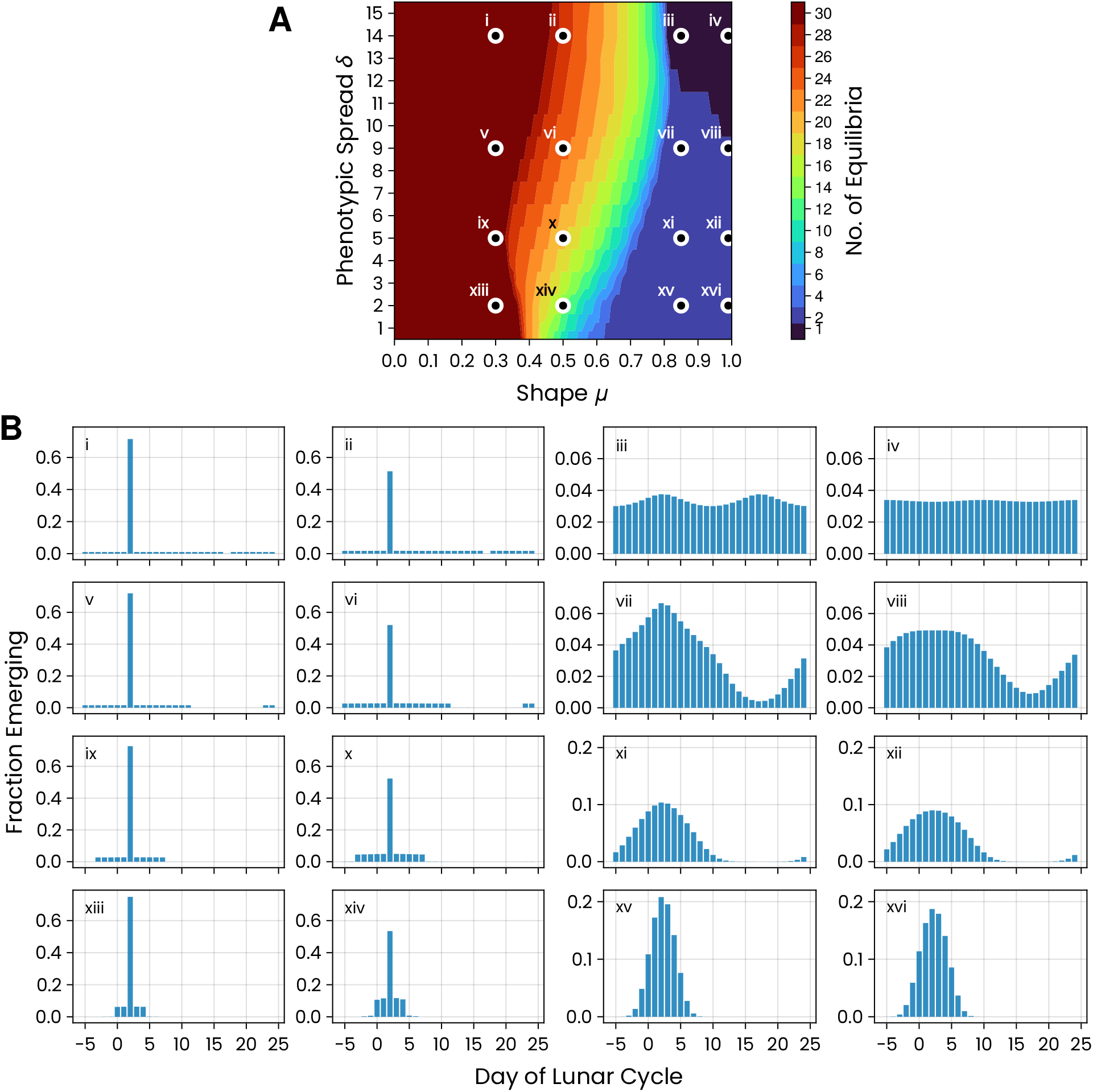
A. The phase diagram depicting the number of solutions to the replicator-mutator equation for M=30 and selection strength *w* = 0.75. It is the same plot as Fig. 3A in the main text. The dots mark the points from which a solution is plotting below. B. The solutions to replicator-mutator equation for different values of shape *µ* and spread *δ* of phenotypic variation as marked and labeled in the contour plots above. The grid of points corresponds to the grid of plots. Only the highest fitness solution is plotted for each different parameter values. The other solutions for a given parameter combination have the same distribution but only shifted in phase.

**Figure S3:**
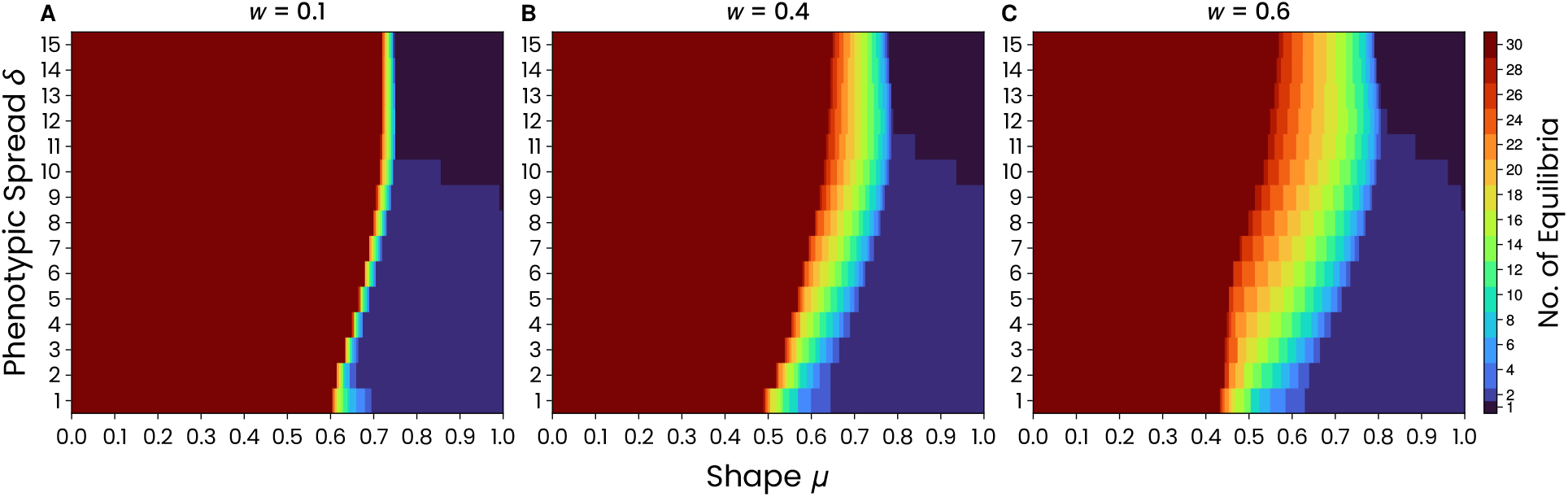
Effect of selection strength on the phase diagram of the parameters of phenotypic variability. A. At low selection strength *w*, the are 30 stable solutions for a large region of the parameter space. Even as the shape parameter *µ* increases until ~ 0.6, these 30 solutions change continuously, but remain stable. But at a certain point, increasing *µ* results in the number of stable solutions decreasing rapidly until we reach the region where lunar and arrhythmic strain are possible (at low and high spread *δ*, respectively). B, C. As *w* increases, the region in which all 30 solutions are stable decreases in size. Solutions with lowest environmental fitness benefit become unstable earlier as *µ* is increased. The region in which it transitions from 30 stable solutions to just 1 or 2 increases in width with increasing *w*. However, the region in which the lunar and arrhythmic strains are seen does not change much in size.

